# Tree nursery environments and their effect on early trait variation

**DOI:** 10.1101/2024.08.20.608769

**Authors:** Annika Perry, Joan K. Beaton, Jenni A. Stockan, Glenn R. Iason, Joan E. Cottrell, Stephen Cavers

## Abstract

Despite the major role of nurseries in raising young plants and trees prior to transplantation, not enough is known about how the nursery climate impacts the growth and development of plants from germination through to maturity. It is important for forestry practitioners to understand the effect that different nursery environments may have on early stage growth as these may exceed differences due to genetic variation and can confound the use of early stage traits for selection. Here, a replicated progeny-provenance experiment of the economically and ecologically important species Scots pine (*Pinus sylvestris* L.) was established in three environmentally distinct nurseries in Scotland and traits including survival, growth, form and phenology were measured. Temperature variation and photoperiod were the only uncontrolled environmental variables during this period, and their effect on measured traits was found to be significant among nurseries from the first growing season onwards. Trait interactions were not consistent between nurseries, indicating that the effectiveness of using proxy traits to select for desirable characteristics may depend on the environment in which the trees are grown. This study is the first in a series that will examine trait variation in Scots pine from seedlings to mature trees and highlights the importance of carefully considering and accounting for the nursery environment when growing trees for subsequent transplantation.

## Introduction

Globally, forests function as the major terrestrial carbon sink (Pan *et al*. 2011) and increasing the area of forest cover is vital to achieving net zero targets (Westaway *et al*. 2023). In addition, increased forest cover would help tackle biodiversity loss, increase filtering of pollutants, provide natural management of flood risk and contribute to soil stabilisation and water purification (Holl and Brancalion 2020). Ongoing afforestation efforts create an increased need for planting stock, which is most commonly supplied as nursery-grown saplings (Brancalion and Holl 2020) and this places the forest nursery in a pivotal and crucially important position. In Great Britain alone, forest nurseries produced around 173 million saplings for the year 2022-2023 (Forestry Commission 2023).

In their role as the primary location for raising young trees, choices made by nurseries have been shown to profoundly influence size and development of young trees, effects that may be observable long after transplantation. For example, the choice of fertiliser (Gruffman *et al*. 2012), growing medium (Heiskanen and Rikala 2000), container type, root management, watering regime (Villar-Salvador *et al*. 1999) and timing of operational activities (such as transplantation, Luoranen 2018) during early life stages can affect root growth, seedling morphology, and susceptibility to pests and pathogens (Selander and Immonen 1992). Whilst soil, water and nutrients can be controlled quite precisely by nursery managers, the geographic location of a nursery and any protection measures used, such as glasshouse cover, determine other critical components of the nursery environment. As variables like temperature and photoperiod can strongly affect the growth, development and biochemistry of seedlings (Downs and Borthwick 1956; Vapaavuori, Rikala and Ryyppö 1992; Nerg *et al*. 1994; Domisch, Finér and Lehto 2001) with effects lasting well beyond transplantation (Dhar *et al*. 2015), the nursery environment can be hugely influential.

The use of locally sourced seed is an explicit requirement when planting native species in the UK (Herbert, Samuel and Patterson 1999), but the location of the nursery in which the resulting plants are raised is rarely specified. For example, the contemporary guiding document for UK forestry – The UK Forestry Standard (Forest Research 2023) - does not comment on nursery management, nor does it connect nursery practice to its recommendations on seed sourcing. Previous publications, such as the Forestry Nursery Practice (Aldhous and Mason 1994), do consider site selection factors, but largely from the point of view of maximising plant performance, minimising risk and logistical considerations. In practice, choice of nursery is generally governed by what the market offers and by economics. Consequently, the nursery site may be environmentally distant from both the site of seed origin and planting site of raised seedlings. If the effect of the nursery environment on plant phenotype is large, then the effect of local seed sourcing may be minimal in comparison. A quantitative evaluation of the impact of the nursery environment on trait variance is therefore potentially very important to seed sourcing policy, nursery practice and to the long-term success of planting initiatives.

The impact of the early growing environment is also a critical factor to consider in tree improvement and breeding programmes, particularly with regards to its effect on survival and on variation in desirable traits. Traditional breeding practice, which makes selections at around half rotation length (Zobel and Talbert 1984), is highly inefficient for trees with long rotation times and there is much interest in methods for screening trait variation at the nursery stage (Wu *et al*. 1997). However, trait values measured prior to transplantation (ca. two years) rarely predict values in later years (Burdon, Bannister and Low 1992; Hong, Fries and Wu 2015; Dong *et al*. 2018) (apart from species with very short rotation lengths, e.g. *Eucalyptus grandis* in Colombia, (Osorio, White and Huber 2003)). The use of ‘proxy’ traits to enable early selection relies on the stability of trait-trait relationships among environments, which may (Fukatsu *et al*. 2011) or may not be (Marron, Dillen and Ceulemans 2007) identifiable. Significant variation in age-age and trait-trait correlations among environments (Williams 1988; Osorio, White and Huber 2003) means that nursery effects may confound selection based on early evaluation of trait variation.

Identifying such nursery effects is particularly relevant for widely planted, economically relevant species. *Pinus sylvestris* L. (Scots pine) is a globally important tree species with a natural range that extends from southern Spain to northern Finland, and from western Scotland to far-eastern Russia. As a foundation species with high conservation value, it has been shown to exhibit high adaptive trait variation at both regional and international scales (Salmela *et al*. 2011; Perry *et al*. 2016; Donnelly *et al*. 2018; Benavides *et al*. 2021; Ramírez-Valiente *et al*. 2021). *Pinus sylvestris* is also an economically valuable timber production species and is widely planted both within and beyond its natural range: it is estimated that it occupies more than 20 percent of the productive forest area in the EU (Mason and Alía 2000) and, within Great Britain, it is the second most abundantly grown species in tree nurseries after Sitka spruce (*Picea sitchensis*, Forestry Commission 2023). Globally, *Pinus sylvestris* grows (both naturally and planted) across a vast range. Its distribution is extremely environmentally heterogeneous and the species has adapted to survive across a massive spectrum of temperature profiles, from hugely variable to relatively stable. *Pinus sylvestris*’ ability to grow across a huge environmental gradient is also associated with a strong phenotypic variation.

Common garden trials are widely used in forestry to quantify the genetic component of phenotypic variation. Replicated common gardens in different environments can be used to examine the environmental component of phenotypic variation. Nurseries, in their efforts to minimise environmental variation within a site and to standardise conditions for each plant, are functionally common gardens. By making use of a controlled, replicated common garden experimental design in nursery settings, we can identify the contribution of the nursery environment to phenotypic variation. This allows evaluation of the importance of the nursery environment to longer term development of the plant and establishes a baseline for interpreting field trial results in years to come. In addition, these results will enable age-age correlations to be explored as the trees mature in field trials.

For this study, we compared early life stage traits of *P. sylvestris* seedlings grown in three nursery environments (subsequently planted in multisite common garden field trials in 2012, for full description see Beaton et al., 2022; Perry et al., 2024b). The plants in each nursery trial had a common genetic background (seed collected systematically from across the natural range in Scotland and arranged in a fully replicated design), were grown under standard soil, nutrient and watering regimes and were managed and measured by the same team of people. As the first paper in a series exploring adaptive trait variation in these trials we focus here exclusively on the effect of nursery environment on overall trait variation (although the experimental design allows for genetic effects on early trait variation to be examined, we do not deal with these here). We quantified the environmental effect on early growth and development and survival and examined relationships within and among traits.

## Methods

### Source of material

Seedling source and nursery environments are detailed in Beaton *et al*. (2022). Briefly, seed was collected from 10 mother trees from each of 21 provenances of *P. sylvestris* from across the native range in Scotland in March 2007. Following stratification, seeds were germinated at the James Hutton Institute, Aberdeen (latitude 57.133214, longitude -2.158764) in June 2007 and transplanted into individual pots. The full final collection consisted of 210 families (10 families from each of 21 provenances) each consisting of 24 half-sibling progeny (total 5,040 individuals).

### Nursery environment

After transfer into individual pots, and with the intention of eventually using these plants to establish field based provenance trials, 8 seedlings per family were moved to one of three nurseries (total 1,680 seedlings per nursery) in July 2007: an unheated nursery glasshouse (NG) at the James Hutton Institute, Aberdeen; an outdoor nursery in the west of Scotland (NW) at Inverewe Gardens, (latitude 57.775714, longitude -5.597181); an outdoor nursery in the east of Scotland (NE) at the James Hutton Institute (location as above). In each nursery, trees were arranged in 40 randomised complete blocks, where each block contained two trees per family (total 42 trees). Watering was automatic in NG, and manually, as required, for NE and NW. The watering regime ensured that water was never limiting and waterlogging of seedlings was avoided. No artificial light was provided in any of the nurseries. In May 2010, the pots containing the seedlings from NG were moved outdoors to Glensaugh in Aberdeenshire (latitude 56.89, longitude -2.54).

Abiotic variation among nurseries was controlled as much as possible by growing the seedlings under standard soil, nutrient and watering regimes. The major sources of abiotic variation which could not be/were not controlled were temperature and wind, of which temperature was measured throughout their growth in the nursery environments. Biotic interactions were not controlled or measured during this period.

Hourly temperature was recorded at each of the nursery sites using data loggers from July 2007 until December 2010. Temperature data loggers were suspended 50 cm above the ground under an aluminium foil-covered funnel. Daily minimum, maximum and average temperatures (between 0000-2300 each day) and the daily temperature variances were calculated for each nursery over this period (Perry et al., 2024a). Daily variance was also partitioned into separate values for the day and night period, where day variance was estimated between the hours of sunrise and sunset between 0000-2300 and night variance was estimated between the hours of sunset and sunrise between 0000-2300. Sunset and sunrise for each day at each nursery were obtained from www.timeanddate.com.

On the few occasions on which more than one hourly record per 24 hours was missing, the daily temperatures were replaced with average daily temperature data derived from the nearest weather stations to the nurseries (information provided by the National Meteorological Library and Archive, Met Office, UK) for the period July 2007 to December 2010). Weather stations were: Poolewe (0.7 miles from NW); Craibstone (4.3 miles from NE and NG) and Glensaugh No 2 (0.1 miles from NG when moved outdoors to Glensaugh). Where weather station data were missing (only relevant for Poolewe over this period) they were replaced with measurements from the next closest weather station: Aultbea No 2 (4.2 miles from NW). Due to the differences in the location between the weather stations and the nurseries and the different methods of recording temperature, data from weather stations were adjusted prior to inclusion in the ‘nursery temperatures’ dataset by comparing daily temperatures across the whole period. Missing logger temperatures were directly replaced, where possible, by checking the weather station temperature at the missing time point and calculating the mean logger temperature recorded at all other instances that the same weather station temperature was recorded. In the few instances where there were no direct replacements, replacement logger values were obtained by taking the mean value for the three logger records either side of the missing logger data point in a dataset which was ordered by the weather station temperatures. For missing data in NG (prior to May 2010), replacement temperatures derived from NE were used instead of weather station data. Where there were days with missing hourly data for one of more of the nurseries, temperature variance could not be estimated and so these days were excluded from analyses for all nurseries to avoid skewing results.

The growing season length in each year at each nursery was estimated as the period bounded by a daily mean temperature >5 °C for >5 consecutive days and daily mean temperature <5 °C for >5 consecutive days (after 1st July). During the defined growing season period trees were assumed to be ‘growing’ whereas outwith the defined growing season trees were assumed to be ‘not growing’. The previously described temperature variances were compared between these two different periods. The growing degrees were estimated for each day in the period July 2007 to December 2010 as the number of degrees above 5 °C. The cumulative growing degree days (GDD) were estimated by summing the growing degrees for each period leading up to phenology assessments at each nursery from 1 January in 2008. Chilling days were defined as the days in which the average maximum temperature did not exceed 5 °C.

### Phenotype assessments

Seedling phenotypes were assessed for all individuals for a range of traits reflecting different life history characteristics: growth, form, phenology and survival.

#### i. Growth traits

Growth traits were assessed annually from 2007-2010. Annual increments in height (HI, mm) and basal stem diameter (DI, mm) were measured as the increase from the end of one growing season to the end of the next (for comparisons of height increment across multiple years, height in the first year of growth in 2007 is considered as the height increment for this year), whilst absolute height (HA, mm) and absolute basal stem diameter (measured at the root collar: DA, mm) were measured at the end of each growing season. Relative growth rate (HR) was estimated as the percentage increase in HI relative to HA in the previous year and was recorded for 2008 to 2010.

#### ii. Form traits

Form was measured annually from 2007-2009. Needle length (NL, mm) was measured for three randomly selected needles per tree and a mean value was obtained. The total number of buds (Bu) on each seedling was counted in 2008 and 2009. Slenderness (HD, the ratio of HA to DA) was recorded in 2010 as a measure of the impact of multiple years of growth on tree form.

#### iii. Phenology traits

Phenology was assessed weekly during the spring and autumn of 2008. Budburst timing (BB) was defined as the number of days from 31 March 2008 to the time when newly emerged green needles were observed. The estimated proportion of trees which had burst bud in each nursery was calculated at each assessment. To compare budburst progression across the three sites, the point at which an estimated 50 % of trees had undergone budburst (BB_50_) was calculated as follows:

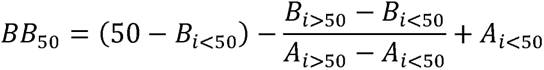

where *i<50* is the *i*th assessment performed at each nursery where < 50 % of trees had undergone budburst (*i<50*: NE = 2; NG = 3; NW = 2); *i>50* is the *i*th assessment where > 50 % of trees had undergone budburst (*i>50*: NE = 3; NG = 4; NW = 3); *A* is the variable associated with each assessment performed at each nursery (*A*: number of days after 31 March; or cumulative growing degree days since 1 January); and *B* is the percentage of trees observed with budburst at each nursery at each assessment.

Growth cessation (GC) was defined as the number of days from 10 September 2008 to the date when no further height growth was observed by eye. To compare the growing season length at each nursery with the duration of growth observed in the seedlings, ‘growth duration’ (GD) was calculated for each seedling as the number of days between budburst and growth cessation.

#### iv. Survival traits

Mortality (number of trees which died by the end of a given growing season as a percentage of the total number of trees alive at the end of the previous growing season) was recorded each year from 2007 to 2010. Survival was recorded as the percentage of surviving trees in each nursery by 2010).

### Statistical analysis

Paired t tests were performed using R (R Core Team 2024) to compare nursery means for annual climatic variables (mean daily temperature, daily variance, growing degree days) and climatic variables measured within and outside the growing season (day temperature variance and night temperature variance). Where there were days with missing hourly data for one of more of the nurseries, temperature variance could not be estimated and so these days were excluded from analyses for all nurseries to avoid skewing results.

Coefficient of variation, calculated as standard deviation divided by the mean, was estimated for each trait in each nursery (Figure S1). Where traits were measured in multiple years only the most recent year was used.

To evaluate relationships among traits, Pearson’s correlation coefficients and significance values were estimated for trees growing in each nursery using the ‘Hmisc’ package (Harrell Jr 2024) in R (R Core Team 2024). Absolute height, absolute basal stem diameter and slenderness were only included for the final assessment year (2010).

Nested analyses of variance (ANOVA) were performed in Minitab version 21 (Minitab 2024) for all trees with nursery as a fixed effect and block nested within nursery as a random effect. Nested ANOVA was also used to analyse the distribution of variance among these traits. In order to account for variance due to relatedness and/or phenotypic plasticity which would otherwise fall within the residual variance, additional terms (provenance; family nested within provenance; a nursery and provenance interaction term) were included but were not considered separately, given that the focus of this study is on the impact of environment on trait variation. Tukey’s post-hoc tests were performed to determine whether mean values among nurseries were significantly different from one another for all traits and for mean temperature, temperature variance (daily, day and night) and growing degree days for each year. ANOVAs were repeated with all terms included as random effects to identify their contribution to the total variance.

## Results

### Climatic variation among nurseries

Trees growing at each of the three nurseries experienced broadly similar mean daily temperatures for each year between 2007 and 2010 which were not significantly different among nurseries (Figure 1) with the exception of NE and NG in both 2008 and 2009, where mean daily temperature at NG was significantly warmer than at NE in both years and warmer in NG than NW in 2009. Trees at NG also experienced the fewest chilling days (total 119 between 2007 and 2010, compared with NE = 157 and NW = 123) and had higher daily GDD throughout the majority of the year (Figure 1) compared to the other two nurseries. In contrast, the daily temperature variance that trees were exposed to was consistently and significantly lower for trees growing at NW than for trees growing at either NG or NE (Figure 1). The daily temperature variance that trees experienced in NG and NE was also significantly different, although the magnitude of the difference was less than when comparing NW with either NE or NG (Figure 1).

**Figure 1.**
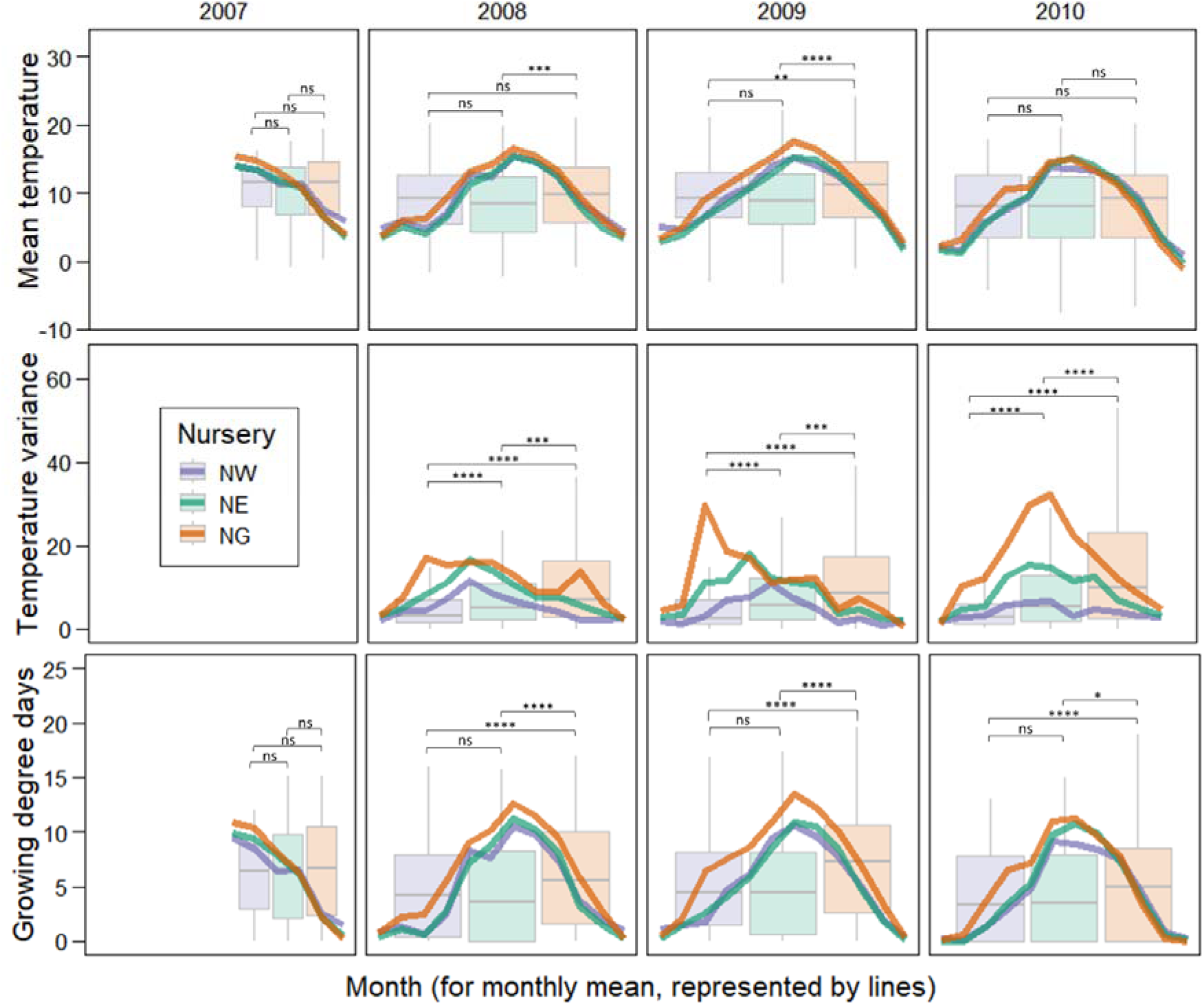
Climatic variation at the three nurseries for the period July 2007 to December 2010 when seedlings were growing in individual pots at the nursery sites. Lines indicate smoothed monthly means for each year. Boxplots indicate values for the entire year: solid grey lines indicate the median value; the bottom and top of the boxes indicate the first and third quartile; and the upper and lower whiskers extend to the highest and lowest values within 1.5 times for interquartile range (no outliers are shown for ease of visualisation. No data are included for daily temperature variance in 2007 due to the extent of missing hourly data for this year. Significant differences among nurseries within each year are indicated with asterisks (*: p = 0.01-0.05; **: p = 0.001-0.01; ***: p = 0.0001-0.001; ****: p < 0.0001; ns: not significant. Units: mean temperature, °C; daily temperature variance, °C^2^, growing degree days, number of degrees above 5 °C for each day.

Differences in temperature variance were most pronounced among nurseries and consistent among years during the daytime within the growing season period (Figure S2). During this period, daytime variance was significantly higher at NG than at NE or NW, and higher at NE than at NW in all years. In contrast, variance during the night-time was lowest at NG compared to the outdoor nurseries in all years outside the growing season (mean night temperature variance 2008-2010: NE = 2.40 °C*^2^*; NG = 1.35 *°*C*^2^*; NW = 2.17 *°*C*^2^*).

### Trait variation among nurseries

Trees growing in NG had higher mean values for traits relating to growth than those growing outdoors in NE and NW by 2010 (Table 1): they had thicker stems and grew taller and faster, on average. They also had larger increments for both height and basal stem diameter in 2010. Trees grown in NE were the smallest and thinnest with the slowest growth rate and lowest increments on average in 2010. Mean values for traits relating to form and survival were lower, and budburst timing was earlier, for trees growing in NG than those growing in one or both outdoor nurseries (Table 1) with the exception of needle length.

**Table 1.** Traits measured in individual seedlings, grouped into the following categories: ‘Growth’, ‘Form’, ‘Phenology’ and ‘Survival’. Overall means and associated standard errors are provided for each nursery (NE, NG and NW) separately. Tukey pairwise comparisons are reported as a subscript letter (A, B, C) after the mean value: nurseries that do not share a letter (within traits for each year) have significantly different means. Number of observations for each nursery are in parentheses.

| Trait/year code* | Unit | NE | NG | NW |
| --- | --- | --- | --- | --- |
| *Growth* |  |  |  |  |
| DA08 | Mm | 3.58<sup>A</sup> ± 0.02 (1465) | 2.87<sup>C</sup> ± 0.02 (1348) | 2.96<sup>B</sup> ± 0.03 (1121) |
| DA09 | Mm | 5.74<sup>A</sup> ± 0.02 (1438) | 4.80<sup>B</sup> ± 0.02 (1248) | 4.69<sup>C</sup> ± 0.04 (1050) |
| DA10 | Mm | 6.77<sup>C</sup> ± 0.04 (1419) | 8.81<sup>A</sup> ± 0.04 (1247) | 7.19<sup>B</sup> ± 0.07 (1029) |
| DI09 | Mm | 2.14<sup>A</sup> ± 0.02 (1438) | 1.90<sup>B</sup> ± 0.02 (1247) | 1.71<sup>C</sup> ± 0.03 (1048) |
| DI10 | Mm | 1.08<sup>C</sup> ± 0.02 (1416) | 4.01<sup>A</sup> ± 0.04 (1246) | 2.48<sup>B</sup> ± 0.04 (1029) |
| HA08 | Mm | 81.12<sup>C</sup> ± 0.5 (1192) | 100.36<sup>A</sup> ± 0.83 (1319) | 88.32<sup>B</sup> ± 0.82 (1014) |
| HA09 | Mm | 237.76<sup>A</sup> ± 1.33 (1440) | 241.14<sup>A</sup> ± 1.9 (1246) | 216.9<sup>B</sup> ± 2.51 (1036) |
| HA10 | Mm | 322.80<sup>C</sup> ± 1.69 (1416) | 382.73<sup>A</sup> ± 2.03 (1245) | 334.87<sup>B</sup> ± 4.13 (775) |
| HI07 | Mm | 59.98<sup>A</sup> ± 0.3 (1611) | 57.30<sup>B</sup> ± 0.32 (1498) | 47.92<sup>C</sup> ± 0.29 (1492) |
| HI08 | Mm | 21.59<sup>C</sup> ± 0.46 (1192) | 42.93<sup>A</sup> ± 0.73 (1319) | 40.14<sup>B</sup> ± 0.80 (1014) |
| HI09 | Mm | 164.83<sup>A</sup> ± 1.23 (1180) | 142.54<sup>B</sup> ± 1.46 (1218) | 138.72<sup>B</sup> ± 2.03 (952) |
| HI10 | Mm | 84.20<sup>C</sup> ± 0.75 (1415) | 141.48<sup>A</sup> ± 1.24 (1238) | 125.84<sup>B</sup> ± 1.85 (767) |
| HR08 | % | 40.11<sup>C</sup> ± 1.08 (1192) | 77.85<sup>B</sup> ± 1.50 (1319) | 90.59<sup>A</sup> ± 2.15 (1014) |
| HR09 | % | 212.06<sup>A</sup> ± 2.05 (1180) | 150.43<sup>C</sup> ± 1.87 (1218) | 162.66<sup>B</sup> ± 2.45 (952) |
| HR10 | % | 36.55<sup>B</sup> ± 0.41 (1415) | 65.81<sup>A</sup> ± 1.07 (1237) | 65.33<sup>A</sup> ± 1.11 (767) |
| *Form* |  |  |  |  |
| Bu08 | Count | 7.14<sup>A</sup> ± 0.07 (1462) | 5.79<sup>B</sup> ± 0.09 (1339) | 4.55<sup>C</sup> ± 0.09 (1118) |
| HD08 | Ratio | 22.56<sup>C</sup> ± 0.16 (1192) | 35.46<sup>A</sup> ± 0.27 (1317) | 29.74<sup>B</sup> ± 0.28 (1014) |
| HD09 | Ratio | 41.67<sup>C</sup> ± 0.21 (1438) | 50.24<sup>A</sup> ± 0.35 (1242) | 45.56<sup>B</sup> ± 0.35 (1036) |
| HD10 | Ratio | 48.50<sup>A</sup> ± 0.25 (1414) | 44.21<sup>B</sup> ± 0.26 (1245) | 47.76<sup>A</sup> ± 0.36 (774) |
| NL07 | Mm | 25.55<sup>B</sup> ± 0.12 (1611) | 26.22<sup>A</sup> ± 0.12 (1495) | 22.84<sup>C</sup> ± 0.13 (1492) |
| NL08 | Mm | 100.42<sup>C</sup> ± 0.36 (1453) | 115.96<sup>A</sup> ± 0.56 (1294) | 104.92<sup>B</sup> ± 0.62 (1113) |
| NL09 | Mm | 31.21<sup>C</sup> ± 0.22 (1440) | 54.67<sup>A</sup> ± 0.39 (1248) | 49.56<sup>B</sup> ± 0.37 (1050) |
| *Phenology* |  |  |  |  |
| BB08 | Days since 31/03/08 | 41.99<sup>B</sup> ± 0.11 (1491) | 19.14<sup>C</sup> ± 0.19 (1454) | 42.58<sup>A</sup> ± 0.16 (1145) |
| GC08 | Days since 10/09/08 | 44.61<sup>B</sup> ± 0.53 (1450) | 44.75<sup>B</sup> ± 0.56 (1393) | 57.61<sup>A</sup> ± 0.35 (1067) |
| GD08 | Days | 165.71<sup>C</sup> ± 0.53 (1440) | 188.76<sup>A</sup> ± 0.57 (1390) | 178.32<sup>B</sup> ± 0.37 (1055) |
| *Survival* |  |  |  |  |
| Su10 | % | 84.88 | 74.23 | 61.31 |
| M07 | % | 4.11 | 10.83 | 10.83 |
| M08 | % | 9.06 | 9.88 | 25.03 |
| M09 | % | 1.64 | 6.44 | 6.41 |
| M10 | % | 1.04 | 1.27 | 2.00 |

\* Trait/year code: Letter(s) denote trait, number denotes year of measurement: traits - DA, absolute basal stem diameter; DI, basal stem diameter increment; HA, absolute height; HI, height increment; HR, relative growth rate; Bu, number of buds; HD, slenderness; NL, needle length; BB, budburst timing; GC, growth cessation; GD, growth duration; Su, survival; M, mortality; years - 07, 2007; 08, 2008; 09, 2009; 10, 2010.

Trait means were significantly different among the three nurseries for each trait and each year with the exception of height increment in 2009 and relative growth rate in 2010, both among NG and NW, and growth cessation among NE and NG (Table 1). The length of needles was considerably longer in all nurseries in 2008 than in either 2007 or 2009 (Figure S3, Table 1). Needles which grew in 2008 were, on average, 3.93 (NE) to 4.59 (NW) times longer than those which grew in 2007, and 2.11 (NW) to 3.21 (NE) times longer than needles which grew in 2009. Overall survival was lower for trees in NW compared to the other two nurseries (Table 1) although this was due to a spike in mortality in 2008 which was thought to be largely due to biotic factors such as sand fleas, snails and fungal infection (although cause of mortality in each case was not recorded). Mortality was highest/joint-highest at NW in three of the four assessment years and was also high in the other year. Mortality at the other outdoor site (NE) was the lowest of all nurseries across all four years. By 2010, there were significant differences among nurseries in seedling survival. Mortality was highest in 2008 at NW (25.03 % annual mortality) which accounted for over half of the seedling deaths across the period for this nursery. By 2010, mortality was low for all nurseries.

Trees that died were those which tended, in the previous year, to be shorter and thinner, formed fewer buds and shorter needles, burst bud and stopped growing later than trees which remained alive (Figure 2). Exceptions to this were found in only six out of 57 comparisons (Figure 2): growth cessation in 2008 for trees growing in NG, some growth traits (absolute and increment height and relative growth rate) for trees growing in NW in 2008 and needle length in 2009 for trees growing in NW and NE.

**Figure 2.**
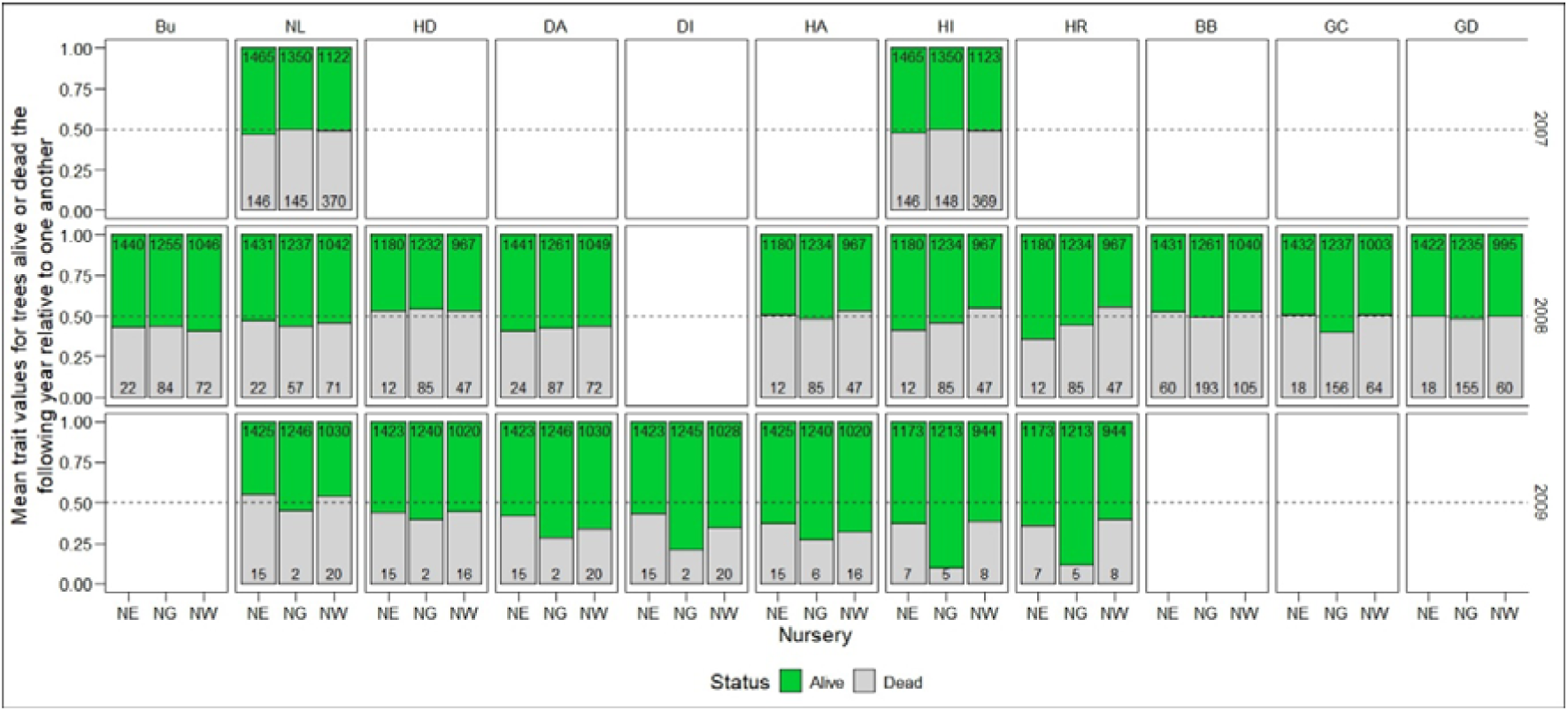
Mean trait values for trees that were either alive or dead by the end of the year after the trait was measured, relative to one another. Where trait means were identical for each group, relative means are each 0.5 (indicated by dashed line). Where means are higher for trees which were alive in the following year, the proportion of the bar that is green exceeds 0.5. Numbers of trees in each group are indicated within the respective coloured section of each bar. Trait codes are given in Table 1.

The relative difference in trait values between trees that were alive and those that died increased as trees got older (Figure 2), although differences over more than one year cannot be compared for traits relating to phenology). The numbers of trees in the ‘dead’ group also reduced over this period, from an average of 221 trees in 2007 to 64 trees in 2008 and 10 trees in 2009 compared to those in the ‘alive’ group (average numbers of trees counted for each trait: 1,313 in 2007; 1,198 in 2008; 1,191 in 2009).

### Associations between nursery climate and trait variation

Timing of phenological traits measured in 2008 was significantly different among nurseries (Table 1): budburst was, on average, 23 days earlier at NG than at either NE or NW; BB_50_ was 38.7 and 38.8 days in NE and NW, respectively, but only 16 days in NG (Figure 3). Over the same period, the difference in GDD among nurseries was much lower (Figure 3, NE: 207.3; NG: 237.7; NW: 238.1). Trees at NE required fewer cumulative GDD, and those at NW and NG required more cumulative GDD to complete budburst throughout the assessment period but only by around 30 degree days. Trees at NW were exposed to two chilling days during the budburst period (5th and 6th April 2008) but the temperature environments were otherwise similar at the two outdoor nurseries (NE and NW, Figure 1).

**Figure 3.**
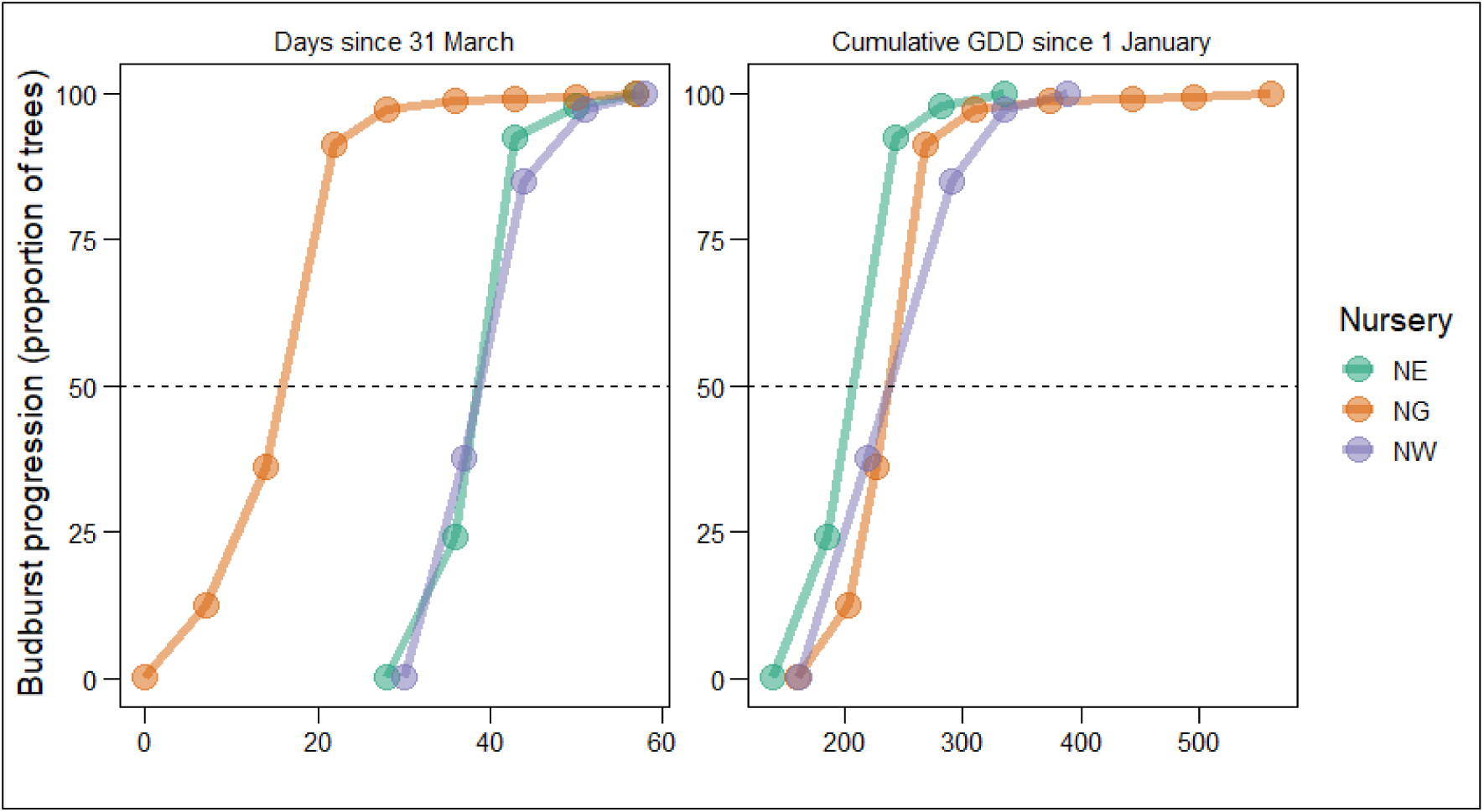
Cumulative percentage of trees reaching budburst, expressed as days since 31 March 2008 (left) and cumulative GDD since 1 January 2008 (right). Circles indicate assessment days. The dotted line indicates the day when 50% of trees had burst bud (BB_50_).

Despite significantly earlier initiation of budburst in NG, trees ceased growing at around the same time as those in NE (Table 1) which was on average about 13 days earlier than in NW. The duration of growth (period between budburst and growth cessation) therefore varied on average between 166 and 189 days (NE and NG, respectively) with trees in NW actively growing for an average of 178 days each year. Mean growth duration was highly significantly positively associated with growing season length (Figure 4a) although the amount of variation in this trait at each site was substantial.

**Figure 4.**
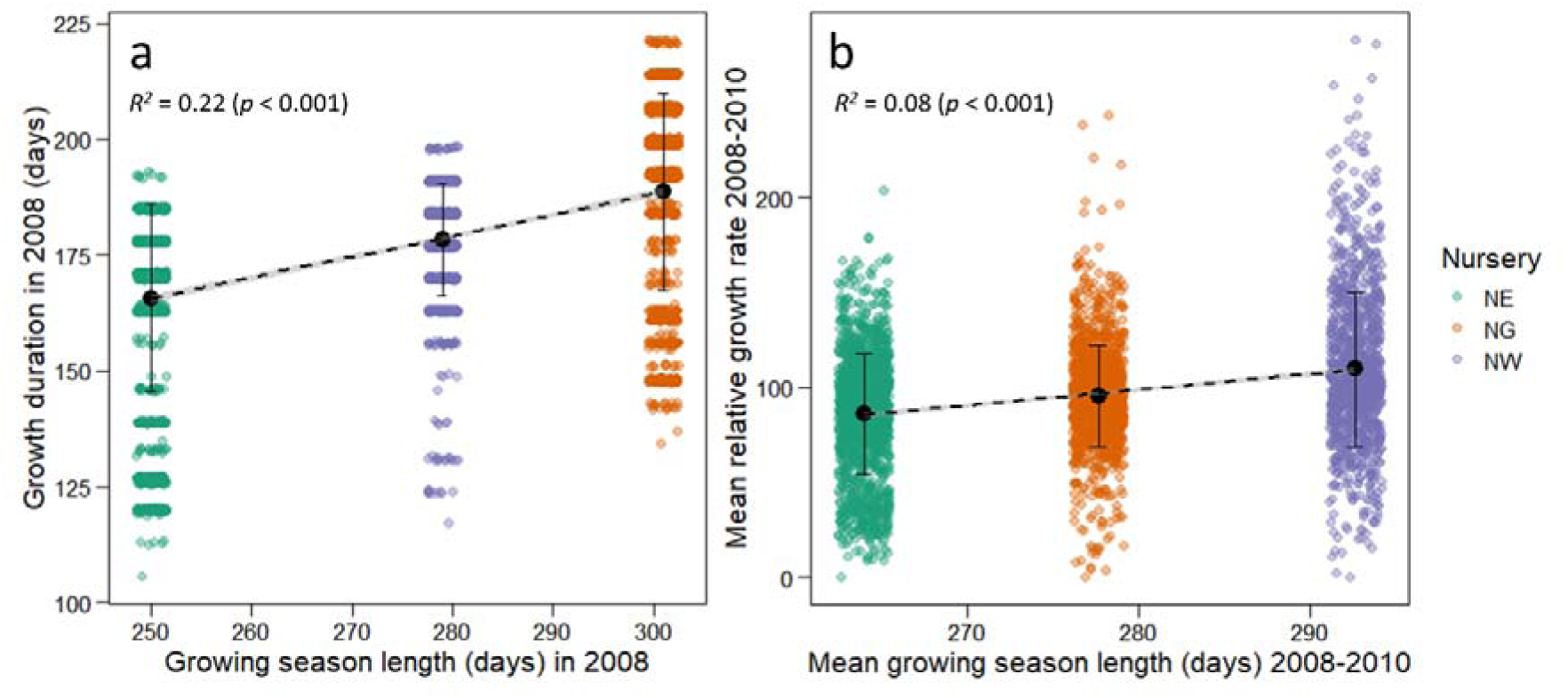
Linear regressions between growing season length (the period in days bounded by a daily mean temperature >5 °C for >5 consecutive days and daily mean temperature <5 °C for >5 consecutive days, after 1st July) and a) growth duration in 2008; and b) relative growth rate 2008 to 2010. Regressions were performed on all data points. Coloured points are trait values for individual seedlings (jittered for clarity). Black points indicate trait mean for each nursery. Error bars are one standard deviation either side of the mean. Mean growth duration: number of days between budburst and growth cessation. Mean relative growth rate: mm per year as a proportion of its size at the end of the previous growing season.

There was also a significant positive relationship between mean relative growth rate and the mean growing season length at each site between 2008 and 2010 (Figure 4b). In-season performance in individual years may be strongly associated with the length of growing season in both the current and the previous year, due to provisioning of the bud during the growing season, but these comparisons will be more powerful using field rather than nursery data, due to the comparatively small number of datapoints available for the latter.

### Relationships within and among traits

Age-age phenotypic correlations were performed for measurements of a given trait obtained in pairs of years for all trees in each nursery site separately (Figure 5). The majority of age-age phenotypic correlations were significant across each pair of years in each of the nurseries, with the exception of needle length (between years 2007 and 2008 in NG; between years 2008 and 2009 in NE), basal stem diameter increment (between years 2009 and 2010 in NG), and height increment (between years 2007 and 2009, and 2007 and 2010 both in NG). Traits were consistently either positively or negatively correlated among years, with the exception of height increment and needle length. Phenotypic correlations among height increments in different years for individuals in NG were negative when 2010 is included (in contrast to the same years in other nurseries for this trait) – this is the year during which the pots were moved outdoors from the glasshouse. The patterns of pairwise comparisons among the sites were more similar for the outdoor sites (NE and NW) than NG. Correlations generally increased in significance as the age of the trees increased and as the number of years between comparisons decreased (Figure 5).

**Figure 5.**
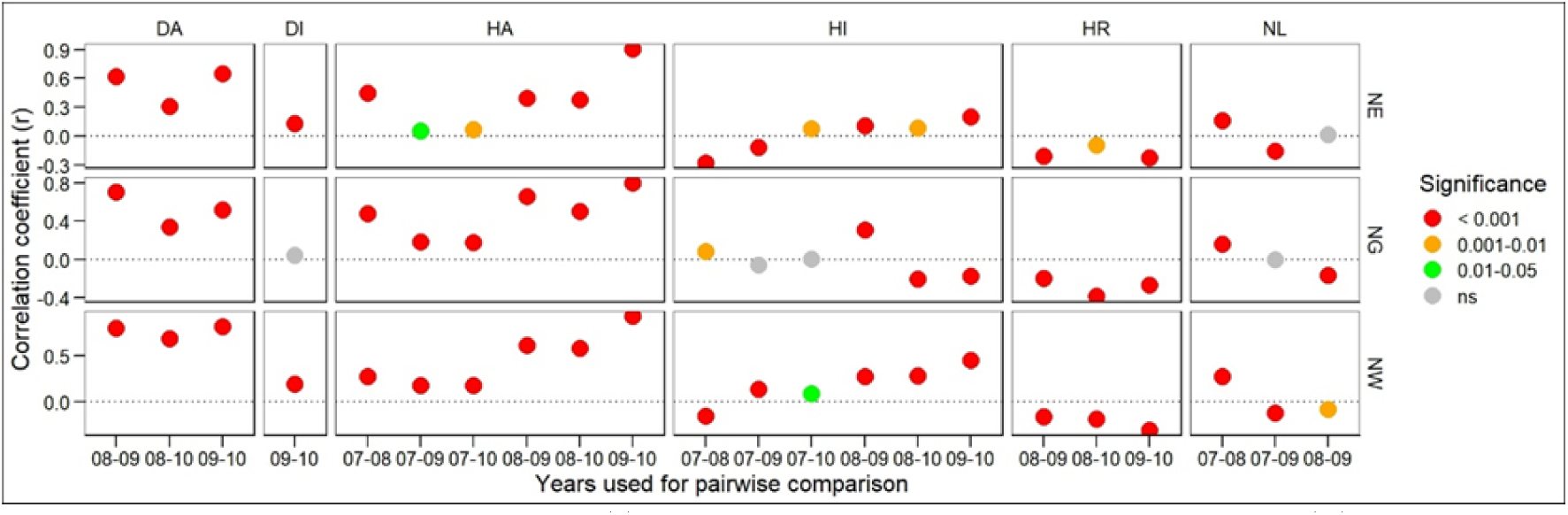
Correlation coefficient (r) and associated significance values (p) for a given trait among years at each nursery (NE, NG, NW). On x-axis, Year is shortened, whereby a pairwise comparison of a trait between years 2007 and 2010 is presented as ‘07-10’. Trait codes are detailed in Table 1. Height measured in the first year (HI07) has been included in both absolute height (HA) and height increment (HI) comparisons

For simplicity, the number of traits measured across multiple years that were consistently significantly associated (in the same direction) among years were reduced prior to subsequent analyses, either to a single (most recent) assessment year (for traits for which successive measurements were not independent: DA, HA) or to a mean value among years (HR, DI). Traits which were not consistent among years (HI, NL) were considered separately for each year.

Pairwise comparisons among growth, form and phenology traits both within and among years (Figure 6) show that pairs of traits were more highly significantly correlated in NW than in either NG or NE, particularly among growth traits and among budburst and other traits. Relationships were often inconsistent among sites: of the 105 pairwise comparisons among traits, 21 were not consistent (i.e. not always either positive or negative, possibly due to the confounding effects of genotype × environment interactions) among nurseries. Of the inconsistent correlations, nearly all (N = 17) involved needle length or height increment (Figure 6). Significantly associated growth traits were always positive, with the exception of height increment in 2007 in NE and NW and in all years at NG (Figures 6).

**Figure 6.**
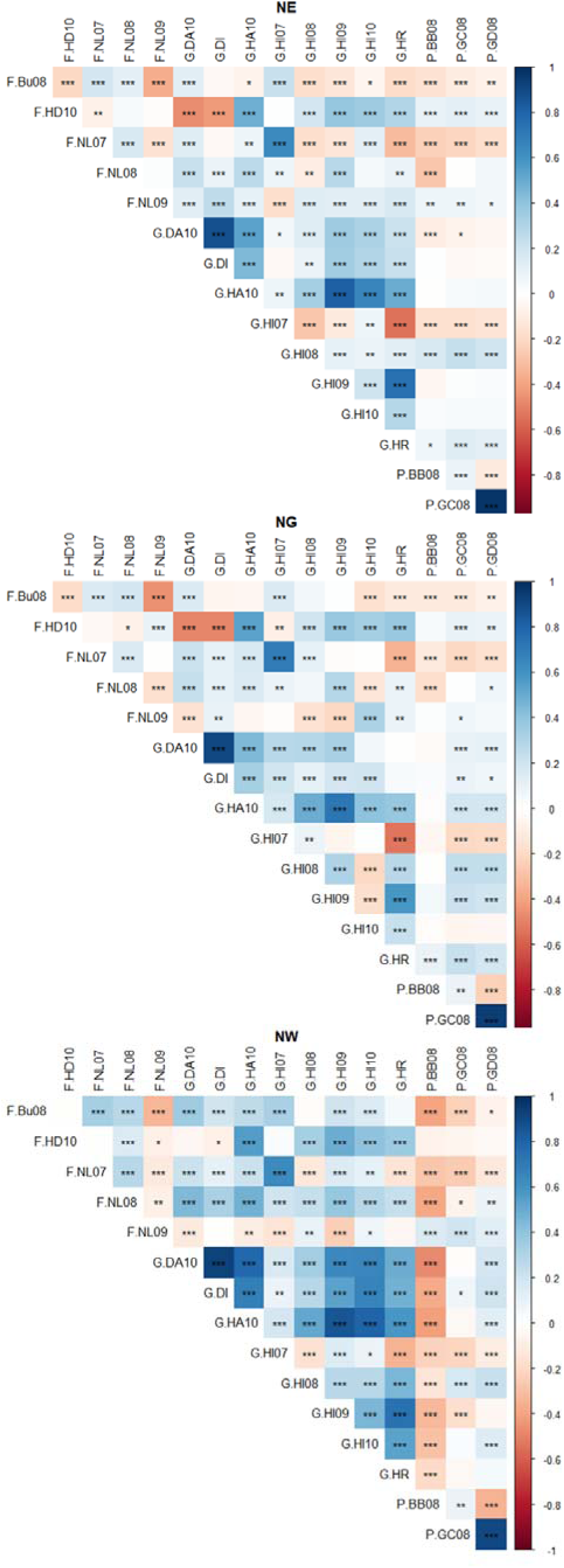
Pairwise correlations for traits for Scots pine trees growing in each nursery (NE, NG, NW); colour indicates value of Pearson correlation coefficient. Significance values: *, p 0.01-0.05; **, p 0.001-0.01; ***, p < 0.001.

Phenology traits were highly significantly positively correlated with one another and were generally negatively correlated with traits related to form (with the exception of slenderness and needle length in 2009). Considering only traits that were measured in the same year (i.e. 2008 for phenology traits), days to budburst were negatively correlated with height increment in NW, positively correlated in NE and not associated in NG. Days to budburst were highly significantly negatively correlated with all growth traits at NW, whereas correlations were either not significant or positively significant for growth traits in NG and NE. In contrast, days to cessation of growth were highly positively correlated with most growth traits in NG (with the exception of height increment in 2007) but associations were much weaker and/or negatively correlated in the other nurseries. Thus, in general, a long growing season between budburst and growth cessation resulted in larger plants (Figure 6).

For further analyses, traits within groups which were consistently highly significantly positively correlated with one another were reduced to a single representative trait (height increment 2008 to 2010, relative growth rate and absolute and increment basal stem diameter were all represented by absolute height in 2010; needle length in 2007 and 2008 were represented by number of buds; growth cessation was represented by budburst)

### Partitioning trait variance

There were highly significant differences among nurseries (i.e. environmental variation over a large scale) as well among blocks within nurseries (environmental variation over a small scale) for every trait (Table S1). Environmental variation (block and nursery) accounted for more than half the total variance for budburst and needle length, close to half for absolute height and around one quarter for the remaining traits (slenderness, number of buds and height increment). Nursery accounted for a greater proportion of variation than block for needle length, number of buds, height increment and budburst (Figure 7), whereas the contribution of block to the total variance was greater than nursery for slenderness and absolute height.

**Figure 7.**
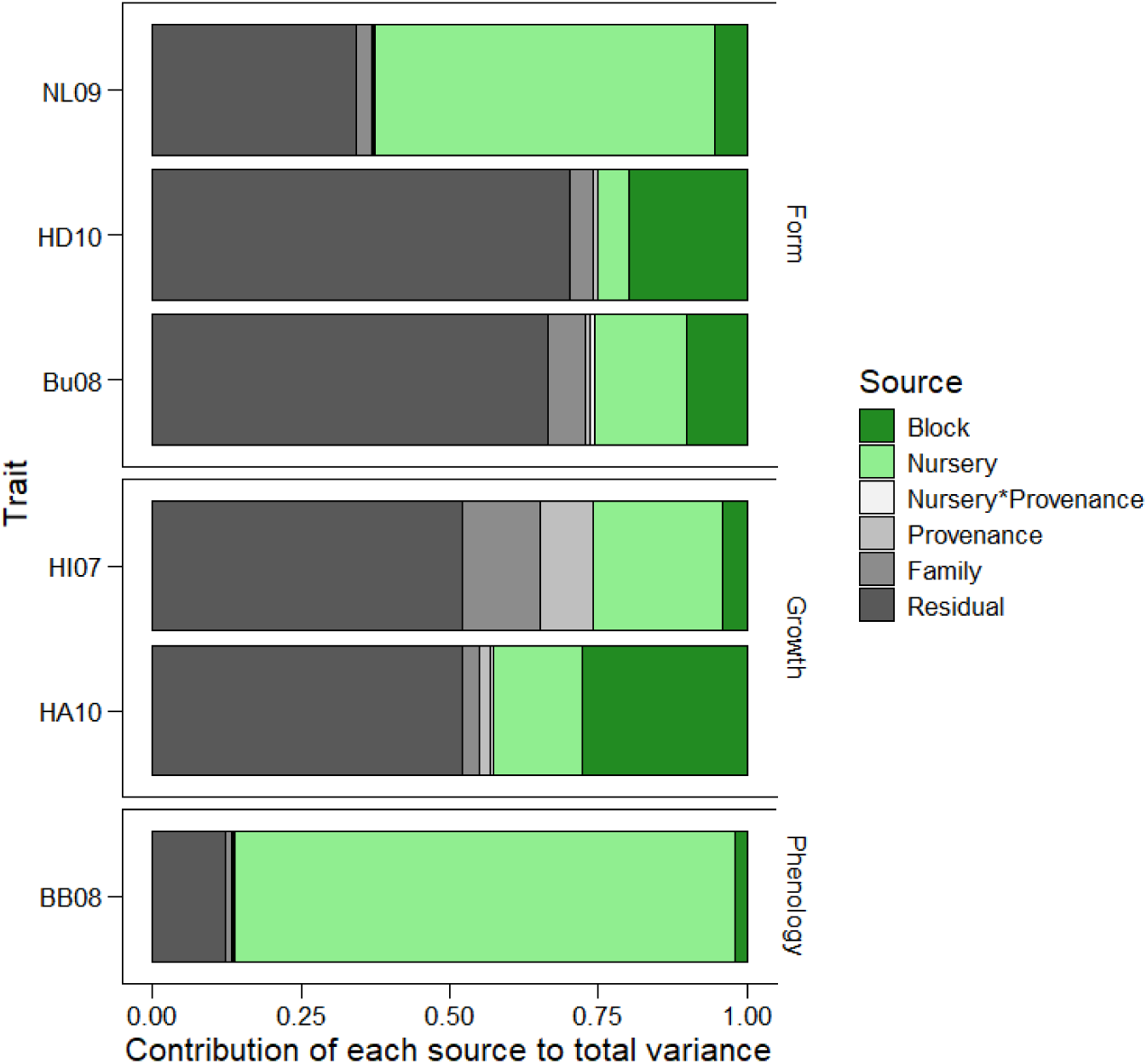
Proportion of variance in growth, form and phenology traits explained by mixed model factors: environmental factors are coloured in shades of green (nursery and block) and genetic/residual factors are coloured in shades of grey (nursery*provenance, provenance, family, residual).

## Discussion

This study highlights the profound impact that nursery environment can have on trait variation in tree seedlings. The nurseries were exposed to the same nutrient, pot and watering regimes, came from the same genetic background and experienced similar average temperatures and monthly growing degree days. Differences in recorded temperature among the nurseries occurred during the daytime within the growing season, when daily temperature variance was much higher for trees growing in the nursery in the east of Scotland compared to the nursery in the west. Seedlings in the glasshouse nursery, in contrast, experienced high daily temperature variance (which was also higher than in the neighbouring outdoor nursery), but temperatures were also higher overall. The importance of the effect of warmer mean temperatures (Morison and Morecroft 2006) and extremes of temperature (Niu *et al*. 2014) on plant traits is increasingly well recognised, especially in the light of an anticipated increase in extreme weather events due to climate change (Herring *et al*. 2022). However, the contrasting effects of temperature variance and mean temperatures and their comparative impact on the growth and development of plants and trees, is less well understood.

In our study, all traits were strongly and significantly affected by the nursery environment: both at a large spatial scale (i.e. among nurseries) but also (and sometimes to a greater extent) at much finer spatial scales (i.e. among blocks within nurseries). The large spatial effect was particularly pronounced between the glasshouse and neighbouring outdoor nursery, highlighting the major impact that nursery type (glasshouse vs outdoor nursery) has on growing plants. While the main environmental difference among the nurseries adjacent (but of different types) to one another was mean temperature, we cannot exclude the possibility that other abiotic (e.g. wind speed, humidity) and/or biotic (e.g. symbionts, pests, pathogens) factors also played a part in the differences we observed.

Growing the seedlings under high temperature variance (resulting in more days at the extreme ends of the temperature range) but with the same mean temperature may lead to lower rates of growth. Chiang, Bånkestad and Hoch (2020) reported significantly taller seedlings when grown in fixed or sinusoidal temperature regimes compared to the same species grown in pots outdoors with a naturally fluctuating and high temperature variance, but only in spring. When grown in the summer, the same seedlings grown in a fixed environment were reported to be significantly shorter than those grown in the outdoor environment, possibly because the lower temperatures that seedlings were exposed to in the outdoor environment in the spring did not occur during summer. The extent of the difference between day time and night time temperatures is also known to directly influence key plant traits relating to growth and form, such as internode length and height (Myster and Moe 1995). Similarly, the allocation of resources to traits relating to tree form in this study appeared to be more important in the nursery with both high variance and low mean temperatures, indicating that a more conservative growth strategy was potentially balanced by investment in traits relating to tree form (Climent *et al*. 2024), such as number of buds.

The role of mean temperature on plant growth is better understood, with higher mean temperatures generally positively associated with higher rates of growth (measured as specific leaf area (Poorter *et al*. 2010). Our study supported this finding, with seedlings grown in a nursery with warmer mean temperatures showing higher rates of growth. Although the temperatures of our nurseries were not controlled, and only a few nursery environments were studied, the environments are analogous to those used in the forestry industry (i.e. outdoor nurseries and glasshouses). We only recorded air temperature, but soil temperature may also have been an important factor for the growth and development of the seedlings. The latter is unlikely to have been consistent among the glasshouse and outdoor environments: it has been reported that soil temperature inside pots tracks but lags behind the air temperature until solar radiation falls directly onto pots, when the temperature of the soil within the pots can rise to more than 20 °C higher than the air temperature (Poorter *et al*. 2016) and that increased soil temperature is associated with increased growth rate (Weih and Karlsson 1999) in saplings.

Growing Scots pine seedlings in nursery environments with different temperature means, variances and photoperiods also had strong effects on phenological variation. Budburst timing was strongly associated with growing degree days at all sites but was not fully explained by differences in this environmental variable among sites. This suggests a complex relationship between warm temperatures (i.e. growing degree days) and cold temperatures (often referred to as the chilling requirement), although the number of nurseries/years assessed in this study is insufficient to enable firm conclusions to be reached on how these two variables interact to result in the observed phenotype. Plant species generally cease growing in response to decreasing photoperiod (Nitsch 1957; Singh *et al*. 2017) although some may respond to decreasing temperature instead (Heide and Prestrud 2005). In our study, trees growing in neighbouring nurseries, outside or within a glasshouse, experienced the same photoperiod (although the light intensity through the glass may have been different) but different mean temperatures. Whereas budburst occurred much earlier in the glasshouse than in the neighbouring outdoor nursery (reflecting the different cumulative growing degree days over the preceding period), growth cessation occurred on average at a similar time which suggests photoperiod to be the main driver.

The accuracy of using young seedlings to predict economic traits at harvest is of major interest and importance in the genetic testing of progeny for forestry (Lee 2002; Hong, Fries and Wu 2015), but detailed early and late assessments of the same tree seedlings are rarely available. Within these experimental trials, which have since been transplanted to field sites in 2012, the same traits will continue to be measured into maturity enabling age-age correlations from seedling to mature trees to be estimated with high resolution (many traits will be measured annually). Although the assessments presented here were confined to the early years of growth this study nevertheless highlights the relative consistency of age-age correlations among the different nursery environments, with the exception of height increment in NG in the years before and after trees were moved outside. The latter showed that plants experienced a shock as a result of the move outdoors which impacted their subsequent growth. The well-documented phenomenon of ‘transplantation shock’ is known to induce changes to growth and development (Close, Beadle and Brown 2005) whereby growth is impaired for a period following transfer. The accuracy of age-age correlations should also, therefore, be reviewed in the context of the general environment in which the trees are measured and the specific environment the trees have been exposed to in the recent past.

The interactions among different traits were also affected by the nursery that plants were grown in. For example, while timing (i.e. lateness) of growth cessation was positively correlated with most growth traits in the protected glasshouse environment, these relationships were weaker in outdoor environments, possibly because these traits were not responding to a single climatic variable but to a combination of many. Similarly, traits relating to form were inconsistent in their relationship with growth traits among nurseries, with more positive associations observed in the nurseries with higher temperature variance compared to the one with low temperature variance.

Given the crucially important role of nurseries in the supply chain for forestry and tree planting, the success of global tree planting efforts depends strongly on the quality of plants they produce. Here we have shown the vital importance of properly understanding the strength and persistence of nursery environment effects on tree seedling growth and development. This knowledge has implications both for genetic testing of material and also for operational tree production. For example, if analyses do not also incorporate data relating to the early environment of the tested trees, an important component of variation might be omitted and may lead to over- or under-estimation of breeding values. Indeed, environmental variation over both small (within sites, measured using blocks) and large (among sites) scales can have significant effects on trait variation and should be carefully recorded and used as covariates in subsequent analyses. For production forestry, although best practice may be to target specific nurseries for the growth of particular species and/or for planting in particular environments (Jaenicke, 1999), economic and practical constraints often render practitioner choice of nursery environment relatively limited.

In this case, improved knowledge of the potential outcomes associated with nursery environment could help practitioners to better understand field traits as an outcome of the environment their trees experienced in the nursery. To this end we recommend routinely monitoring and making available nursery environmental data alongside plant provenance data on shipment from the nursery. The significant effect of nursery environment on all aspects of phenotype reported in this study highlights the cumulative and divergent effect that the early growing environment can have, an effect that may have particularly profound and persistent effects on long-lived species like trees. These findings will be investigated and tested further for the subset of trees that were transplanted to field locations in 2012, using measurements made over many subsequent years.

## Conclusions

Despite the relatively low levels of environmental variation recorded among the three nurseries, there were significant effects on seedling mortality, variation in traits relating to growth, form and phenology and on interactions among traits. These findings have implications for the choice of nursery in which plants are raised given that effects may persist and affect subsequent performance in the field. Growers may wish to assess whether plants should be preferably raised in less benign nursery environments in order to minimise the shock they receive when transplanted. In any case, we are confident of the importance of measuring and collecting data at an early stage in order to compare trait variation throughout the lifetime of trees, from seedling through to maturity: an approach which would be of benefit for plants grown for both research and for operational purposes. The trees in this study have now been transplanted to three field environments (Beaton *et al*. 2022) and subsequent analysis explores the persistence of early environment carry over effects following transplantation (Perry et al., 2024b). Regular measurements on the trees since transplantation will also enable ontogenetic effects on intraspecific genetic variation to be characterised for a wide range of phenotypic traits and for age-age correlations from seedling to maturity to be estimated.

## Acknowledgements

We thank Stewart Mackie and staff at Forest and Land Scotland (Selkirk Office), the Forestry Commission, National Trust for Scotland, and Donald Barrie (Glensaugh) for their help in locating, establishing and securing access to sites.

## Data availability statement

The data underlying this article are available from the Environmental Information Data Centre (EIDC, https://eidc.ac.uk), at https://doi.org/10.5285/29ced467-8e03-4132-83b9-dc2aa50537cd. Temperature and temperature variances at each nursery are available at https://doi.org/10.5285/81841d93-41e2-47a7-b15a-92d1e1cf07f7.

