## Supplementary material for "Tree nursery environments and their effect on early trait variation": Figure S1; Figure S2; Table S1

Supplementary materials

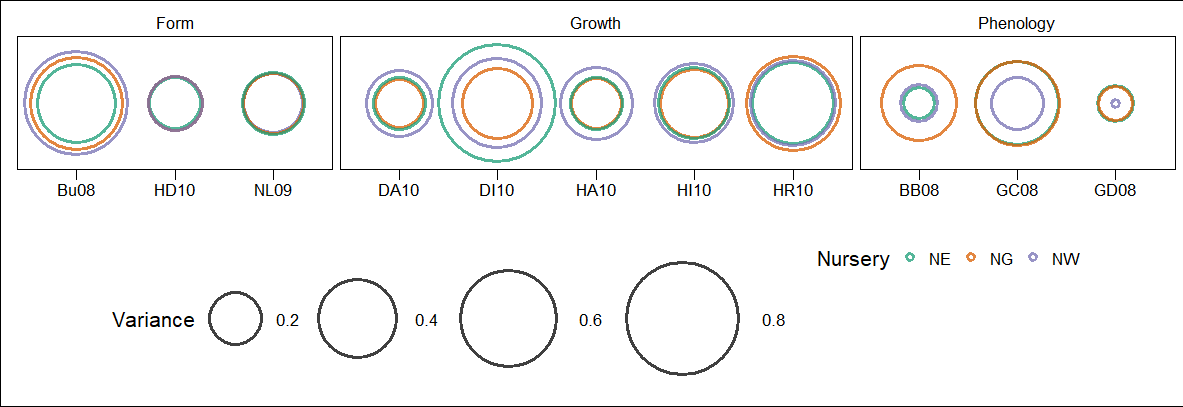

*Figure S1. Coefficient of variation for trait values recorded for Scots pine growing in three nurseries: NE, NG and NW. For traits measured in multiple years, only the most recent year is shown. Coefficients of variation (CV) for each trait at each nursery were generally the inverse of observed mean values (see Table 1), for example: whereas trees growing at NG had the highest mean values for all traits relating to growth, they had the lowest CVs for growth traits, with the exception of relative growth rate.*

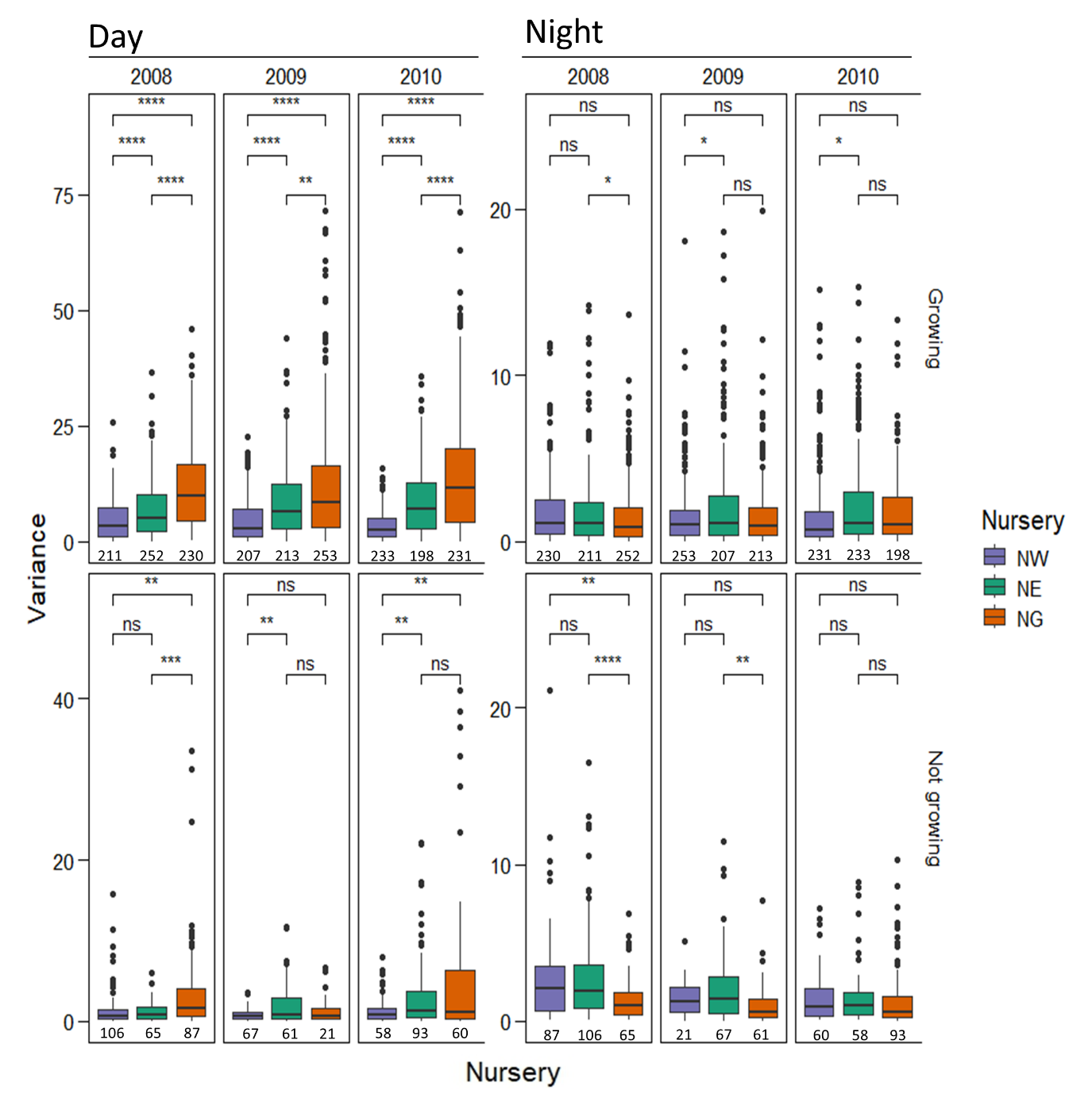

Figure S2. Box and whisker plots showing variation in temperature variance during the night (between sunset and sunrise) and during the day (between sunrise and sunset) both within the growing season (“Growing”) and outside of the growing season (“Not growing”), estimated as the period bounded by a daily mean temperature >5 °C for >5 consecutive days and daily mean temperature <5 °C for >5 consecutive days (after 1st July) for each nursery in each year from 2008 to 2010. No data are included for 2007 due to the extent of missing hourly data for this year. Significant differences among nurseries within each year are indicated with asterisks (*: p = 0.01-0.05; **: p = 0.001-0.01; ***: p = 0.0001-0.001; ****: p < 0.0001; ns: not significant). Boxplots indicate values which fall within the growing season or outwith the growing season for the entire year indicated: solid grey lines indicate the median value; the bottom and top of the boxes indicate the first and third quartile; the upper and lower whiskers extend to the highest and lowest values within 1.5 times for interquartile range; outliers are indicated with black circles. The number of records for each period are indicated under the boxplot.

…………………………………………………………………………………………………..

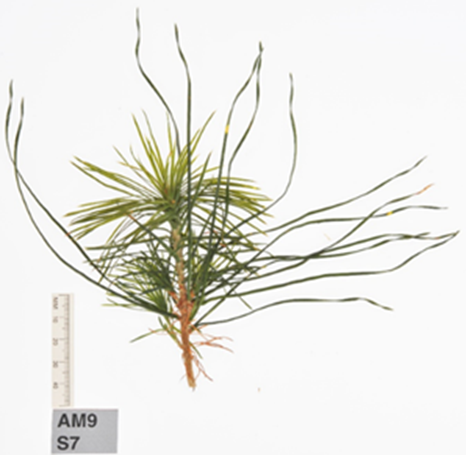

Figure S3. Needle length in all three nursery environments was noticeably greater in the second year of growth (2008) compared to the first and third years (2007 and 2009, respectively).

…………………………………………………………………………………………………..

Table S1. Adjusted mean sum of squares and degrees of freedom (df) from nested analyses of variance for growth and phenology traits (codes given in Table 1) from 2007 to 2010 (07-10) assessments using a subset of traits and years. Parentheses indicate nesting. Significance values: ns, not significant; *, p 0.01-0.05; **, p 0.001-0.01; ***, p < 0.001. Degrees of freedom: block = 117; nursery = 2)

|  | Adjusted mean sum of squares | | | |
| --- | --- | --- | --- | --- |
| Trait/Year | Block (Nursery) | Nursery | Residual | Residual df |
| *Growth* | | | | |
| HA10 | 65896*** | 1220620*** | 4395 | 3316 |
| HI07 | 406*** | 60460*** | 136 | 4481 |
| *Form* | | | | |
| Bu08 | 45*** | 2177*** | 8 | 3799 |
| HD10 | 597*** | 6042*** | 72 | 3313 |
| NL09 | 638*** | 211548*** | 114 | 3618 |
| *Phenology* | | | | |
| BB08 | 181*** | 247368*** | 28 | 3970 |
